# A mathematical framework for human neutrophil state transitions inferred from single-cell RNA sequence data

**DOI:** 10.1101/2025.06.27.662068

**Authors:** Gustaf Wigerblad, Jonathan Carruthers, Sumanta Ray, Thomas Finnie, Grant Lythe, Carmen Molina-París, Saumyadipta Pyne, Mariana J. Kaplan

**Author notes:** These authors contributed equally (Gustaf Wigerblad, Jonathan Carruthers, Carmen Molina-París, Saumyadipta Pyne, Mariana J. Kaplan).

## Abstract

Neutrophils, the most abundant immune cells in the human circulation, play a central role in the innate immune system. While neutrophil heterogeneity is a topic of increasing research interest, few efforts have been made to model the dynamics of neutrophil population subsets. We develop a mathematical model to describe the dynamics that characterizes the states and transitions involved in the maturation of human neutrophils. We use single-cell gene expression data to identify five clusters of healthy human neutrophils, and pseudo-time analysis to inform model structure. We find that precursor neutrophils transition into immature neutrophils, which then either transition to an interferon-responsive state or continue to mature through two further states. The key model parameters are the transition rates (the inverse of a transition rate is the mean waiting time in one state before transitioning to another). In this framework, the transition from the precursor to immature state (mean time less than an hour) is more rapid than subsequent transitions (mean times more than 12 hours). Approximately a quarter of neutrophils are estimated to follow the interferon-responsive path; the remainder continue along the standard maturation pathway. We use Bayesian inference to describe the variation, between individuals, in the fraction of cells within each cluster.

## Introduction

Neutrophils are the most abundant immune cell subset in the human circulation and play a central role in the rapid innate immune response against microbes and other danger signals [1]. Neutrophils are endowed with efficient strategies to neutralize micro-organisms, including phagocytosis, degranulation, and the formation of neutrophil extracellular traps (NETs). Conversely, neutrophil dysregulation or enhanced activation can contribute to auto-inflammatory and auto-immune diseases and promote end-organ damage, while decreases in neutrophil function or numbers can lead to enhanced risk for infections [2, 3]. The estimated half-life of human mature neutrophils in circulation is considered to be less than a day, which contributes to the challenges to fully characterize and study them [4].

Indeed, despite being such a crucial component of the innate immune system, neutrophil physiology and dynamics remain less well characterized than what is known for other immune cell types. This is likely due to technical difficulties in the study of neutrophils, stark differences in neutrophil behavior across certain species, and the short life in circulation of these cells. Recent technological advances allow for the investigation of transcriptional states of single neutrophils, which has begun to elucidate their heterogeneity and complexity [5, 4]. We recently showed that neutrophils can be described along a transcriptional continuum, with discrete clusters based on expression of certain genes [6]. However, accurate knowledge of their cell state dynamics remains incomplete, and is important to characterize the functionality of different neutrophil subsets.

Mathematical modeling has been used to describe the growth, differentiation, and decay of immune cell populations. These models are often based on different hypotheses about the biological mechanisms involved [7]. By comparing model predictions with experimental data, researchers cannot only evaluate the plausibility of these hypotheses but also guide the design of future experiments. So far, much of this work has focused on the generation and maintenance of immunological memory, particularly through models of T-cell populations [8]. This includes models of CD8^+^ effector T-cell production, as well as more recent approaches that use single-cell clustering and pseudo-time trajectory analysis to track how tumor cells develop drug resistance [9, 10]. In contrast, relatively little effort has been made to mathematically model the transcriptional changes in neutrophils or to quantify their movement through the body beyond what has been observed *in vitro* [11].

While previous work [6] identified four transcriptionally distinct neutrophil states using single-cell RNA sequencing, it did not provide a quantitative model of their transitions or kinetics. Our study builds on this by introducing a fifth precursor state and developing an integrated mathematical and statistical framework that models the dynamic transitions between neutrophil states. This enables estimation of transition rates, as defined by the mathematical model, and provides a mechanistic interpretation of cell state progression, which was not captured in earlier analyses [6].

## Methods

### Analysis of scRNA-seq data

Human peripheral neutrophils were purified from adult healthy volunteers as described previously [6]. Cells where captured and libraries made using the 10X 3’ 3.1 chemistry. Matrix files were generated using Cell-ranger 8.0.1 for downstream analysis. The Python package scanpy (version 1.9.4) was used to import matrix files. Cells with fewer than 10^2^ genes or a mitochondrial gene proportion greater than 0.1 were removed. Genes that were expressed in fewer than 5 cells were also omitted from the subsequent analyses.

Healthy volunteer samples were integrated using scvi-tools (version 1.3) and the top 3 *×* 10^3^ variable genes. This creates a low-dimensional, latent representation of the raw gene expression data that attempts to capture the true biological variation while correcting for batch effects. The latent representation is then used to construct a neighborhood graph that identifies cells that are similar to each other and forms the basis of the clustering. Embedding of this graph in two dimensions is performed using Uniform Manifold Approximation and Projection (UMAP) and allows for better visualization of the clusters. Initial Leiden clustering at a resolution of 0.5 identified eight clusters. Cells in three of these clusters were removed as contaminants due to low expression of *CSF3R, FCGR3B* and *NAMPT*, genes that are expected to be highly expressed in human neutrophils. The scVI model was retrained on cells in the remaining clusters and a resolution of 0.42 was used to identify five clusters associated with different neutrophil transcriptional states.

### Mathematical model

Let *n*_*i*_(*t*) denote the concentration of circulating neutrophils in state *i* (with 1 ≤ *i* ≤ *N*) at time *t* ≥ 0, where *N* is the number of states identified from the clustering analysis of individual cells. The influx of precursor neutrophils (or precursor state), *n*_0_, is assumed to occur with constant rate *ϕ*, and neutrophils in any state are lost from circulation at rate *µ*. A neutrophil transitions from state *i* to state *j* with rate *ξ*_*i,j*_. The changes in neutrophil populations associated with each state may then be described by the following system of ordinary differential equations (ODEs):

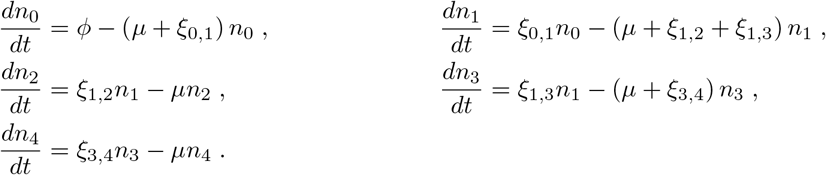

This system reflects our hypothesis that precursor neutrophils, *n*_0_, enter state 1 before differentiating into either state 2 or state 3. The population of neutrophils in state 3 then proceed to the most mature state, represented here by state 4 (see Figure 3).

For healthy individuals, we assume that these populations are in equilibrium (stable steady state), and define *f*_*i*_ to be the fraction of circulating neutrophils in state *i*; that is, *f*_*i*_ = *n*_*i*_*/*∑_*j*_ *n*_*j*_. The transition rates may then be written in terms of these steady state fractions, all of which are measurable, and the loss rate, *µ*, as follows

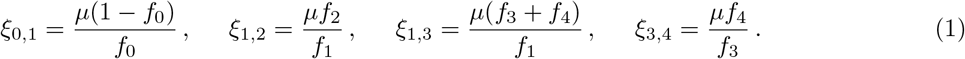

### Statistical model and inference

To understand the variability in transition rates across a population of individuals, we begin by using a Dirichlet distribution, Dir(***α***), to model the population-level variability in steady-state neutrophil fractions. A random variable following a Dirichlet distribution takes values in the form of *N* -dimensional vectors, with the conditions that each entry of the vector is non-negative and all entries sum to one. Since these conditions apply to the steady-state fractions of neutrophils, the Dirichlet distribution provides a natural way to model the between-individual variability in these fractions.

The aim here is, therefore, to estimate the parameters of the Dirichlet distribution, ***α***, using the cluster fractions derived from the analysis of the single-cell sequencing data, our observed fractions. Let ***F*** be a *M × N* matrix of these observed fractions, where *M* = 7 is the number of subjects we have samples for and *N* = 5 is the number of neutrophil states. The *m*^th^ row, ***F*** ^(*m*)^, then contains the cluster fractions for individual *m* (1 ≤ *m* ≤ *M*). For a particular value of ***α***, the likelihood of our observed fractions is given by

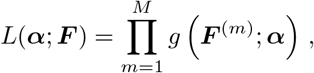

where *g* is the probability density function of the Dirichlet distribution.

The frequentist approach of maximum likelihood estimation looks to find a single ***α*** that maximizes the above likelihood, or equivalently, the value of ***α*** that makes the observed data most probable. Here, however, we adopt a Bayesian approach that treats our parameters as random variables. In this case, the distribution of ***α*** in the presence of our data, or the posterior distribution, is assumed to be proportional to the product of our prior beliefs and the above likelihood [12]. Often, as is the case here, precise expressions for the posterior distribution are difficult to obtain, so computational methods such as Markov Chain Monte Carlo (MCMC) are used to instead sample from it. The posterior sampling is performed here using the emcee Python package, version 3.1.4 [13]. To construct distributions for the transition rates, *ξ*_*i,j*_, ***α*** are first drawn from the MCMC posterior sample. Then, for each draw, steady-state fractions *f*_*i*_, 1 ≤ *i* ≤ *N*, are sampled from the corresponding Dir(***α***) distribution and these sampled fractions are substituted into Equation (1). In this way, we account for both the uncertainty in our estimate of ***α***, and the variability in steady-state fractions across a population of individuals.

## Results

Clustering of neutrophils from seven human samples at single-cell level has led to identification of five clusters corresponding to different transcriptional states. These clusters are depicted in Figure 1, where the UMAP method of dimensionality reduction allows us to visualize the similarity between cells in two dimensions. Full details of the process used to derive these clusters is provided in the Methods section. To interpret the clusters, the average gene expression of cells in each cluster is evaluated for a subset of genes (Figure 2). This subset comprises the top marker genes for each of the clusters found here, as well as marker genes from a previous study of the same dataset [6].

**Figure 1.**
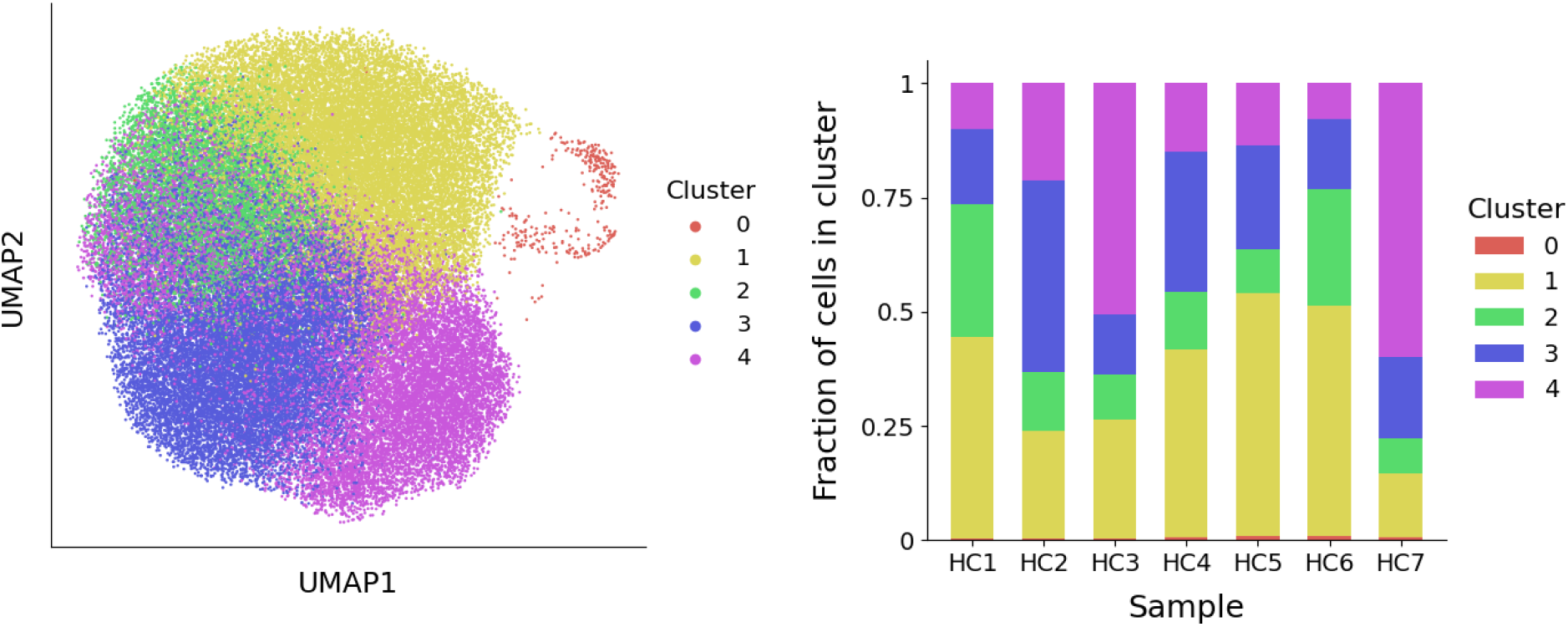
**Left**: A visualization of the five neutrophil clusters that uses UMAP to produce a two-dimensional representation of the raw gene expression data. **Right:** For each healthy subject (HC1-HC7), the fraction of neutrophils that belong to each cluster is depicted.

**Figure 2.**
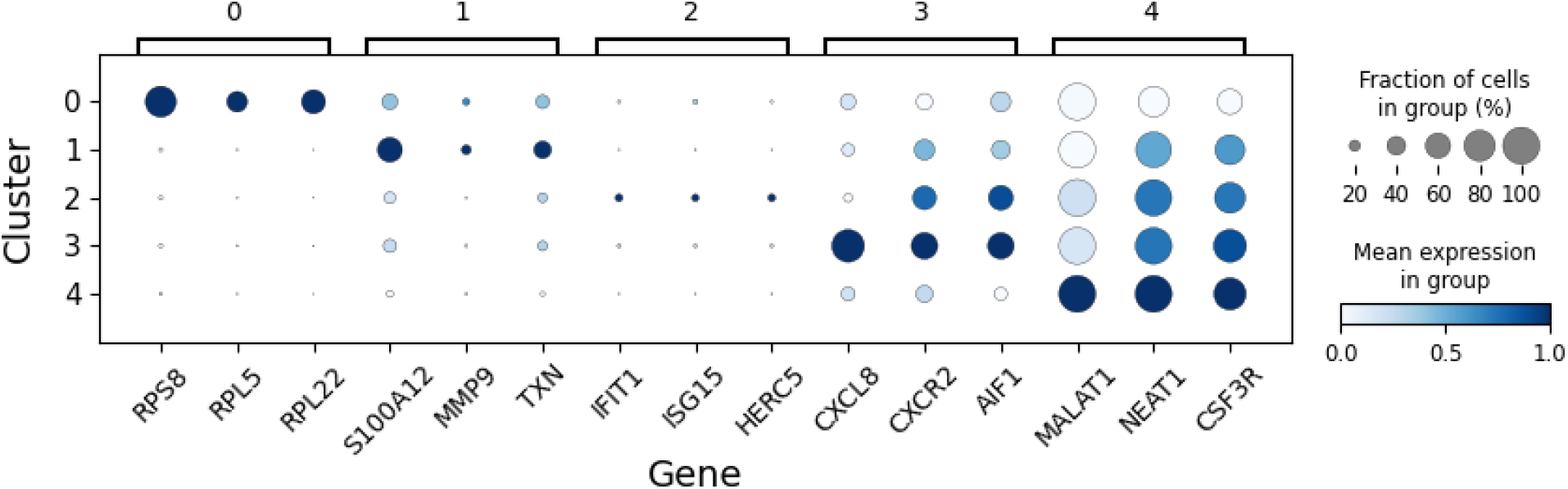
The average expression of select genes in neutrophils in each of the five clusters (states). These expression levels guide the interpretation of each cluster.

Cluster 0 is characterized by elevated expression of ribosomal protein genes (*RPL, RPS*), suggesting these are precursor neutrophils that have recently entered circulation from the bone marrow. Cluster 1 shows higher expression of S100 family genes and likely represents immature neutrophils. Cluster 2 defines a distinct state, marked by up-regulation of genes associated with a type I interferon (IFN) response. Clusters 3 and 4 correspond to increasingly mature circulating neutrophils, distinguished by elevated expression of *CXCR2* and the long non-coding RNAs, *MALAT1* and *NEAT1*, respectively. This interpretation of neutrophil states aligns with our previous study, which identified four transcriptionally distinct clusters corresponding to clusters 1-4 described here [6].

To better understand neutrophil dynamics across the identified clusters, we developed a mathematical model based on a system of ODEs, a standard approach for describing how quantities evolve over time. In this model, each variable represents the concentration of neutrophils in one of the five transcriptional states. The concentration within a state can increase due to incoming cells from other states and decrease due to transitions to other states or exit from circulation. As such, the model requires a defined ordering of states. Previously, pseudo-time analysis was used to order neutrophils along a biological continuum based on gene expression profiles, offering a plausible path through transcriptional states [6]. We used this pseudo-time ordering to inform the structure of our model (illustrated in Figure 3). In this framework, precursor neutrophils (*N*_0_) enter the circulation from the bone marrow and transition into immature neutrophils (*N*_1_).

**Figure 3.**
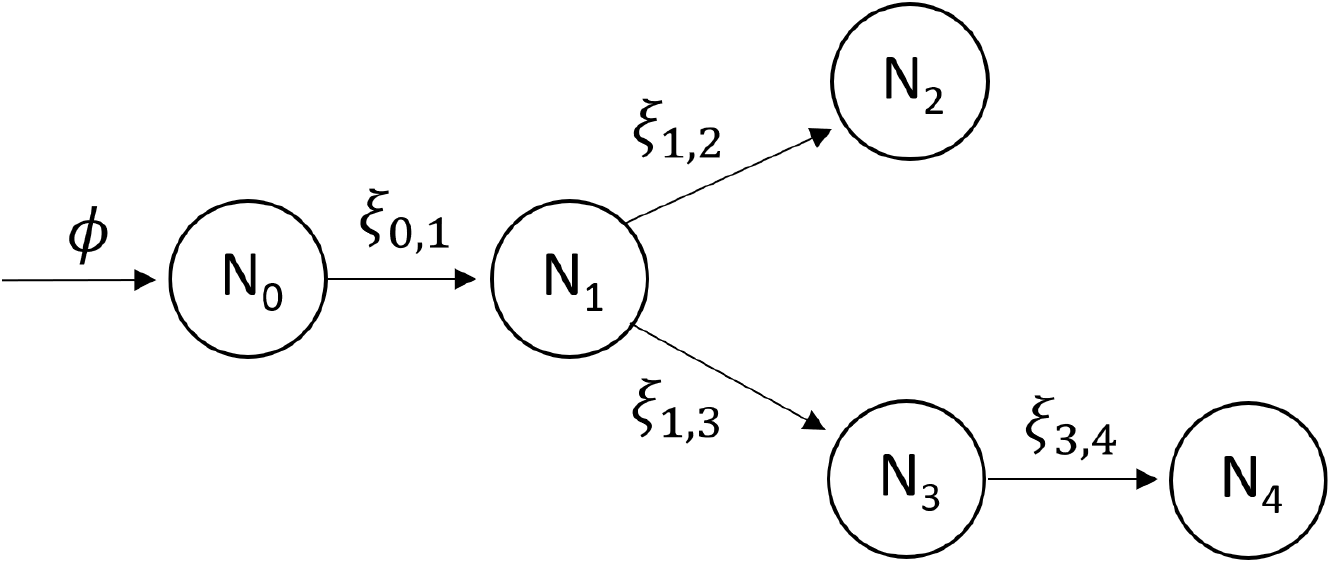
The proposed mathematical model for the transition of neutrophils between different transcriptional states. Neutrophils enter the circulation from the bone marrow at rate *ϕ*, with subsequent transitions between states marked by arrows. The state subscripts correspond to the cluster number and the parameter *ξ*_*i,j*_ represents the rate at which a neutrophil transitions from state *N*_*i*_ to state *N*_*j*_. Neutrophils in each state are cleared at rate *µ*, however, the arrows depicting these events are omitted from the diagram to improve clarity.

From there, cells either pass through an IFN-responsive state (*N*_2_) or continue to mature through states *N*_3_ and *N*_4_. The model’s key parameters are the transition rates, *ξ*_*i,j*_, representing the rate at which neutrophils move from state *N*_*i*_ to state *N*_*j*_. The inverse of these rates (1*/ξ*_*i,j*_) corresponds to the average time spent in state *N*_*i*_ before transitioning to *N*_*j*_. High transition rates imply short dwell times, whereas low rates indicate longer persistence in a state. To estimate these parameters, we assume that neutrophil populations in healthy individuals are at steady-state, meaning the number of cells in each state remains constant over time. Under this assumption, algebraic equations can be derived for the steady-state neutrophil fracions in each cluster, *f*_*i*_, that depend solely on the transition rates and a fixed exit rate from circulation. This exit rate is set based on a presumed 12-hour half-life for circulating neutrophils [14]. We then estimate the transition rates of the seven healthy individuals by substituting their observed cell fractions (Figure 1, right) into these steady-state expressions.

Since cluster fractions vary across individuals, the corresponding estimates of transition rates also differ. However, due to the small sample size, these estimates alone do not capture how transition rates vary across the broader population. To address this, we developed a statistical model, described in detail in the Methods section, that provides us with predicted probability distributions for the steady-state cluster fractions (Figure 4). Using Bayesian inference, we obtain full posterior distributions, where each sample corresponds to a possible realization of cluster fractions. By drawing random samples from this statistical model, we simulate cluster fractions that are representative of a broader population. These samples are then used to infer corresponding probability distributions for the transition rates in the mathematical model (Figure 5). Together, the statistical and mathematical models link individual-level data to population-level estimates of neutrophil state transitions.

**Figure 4.**
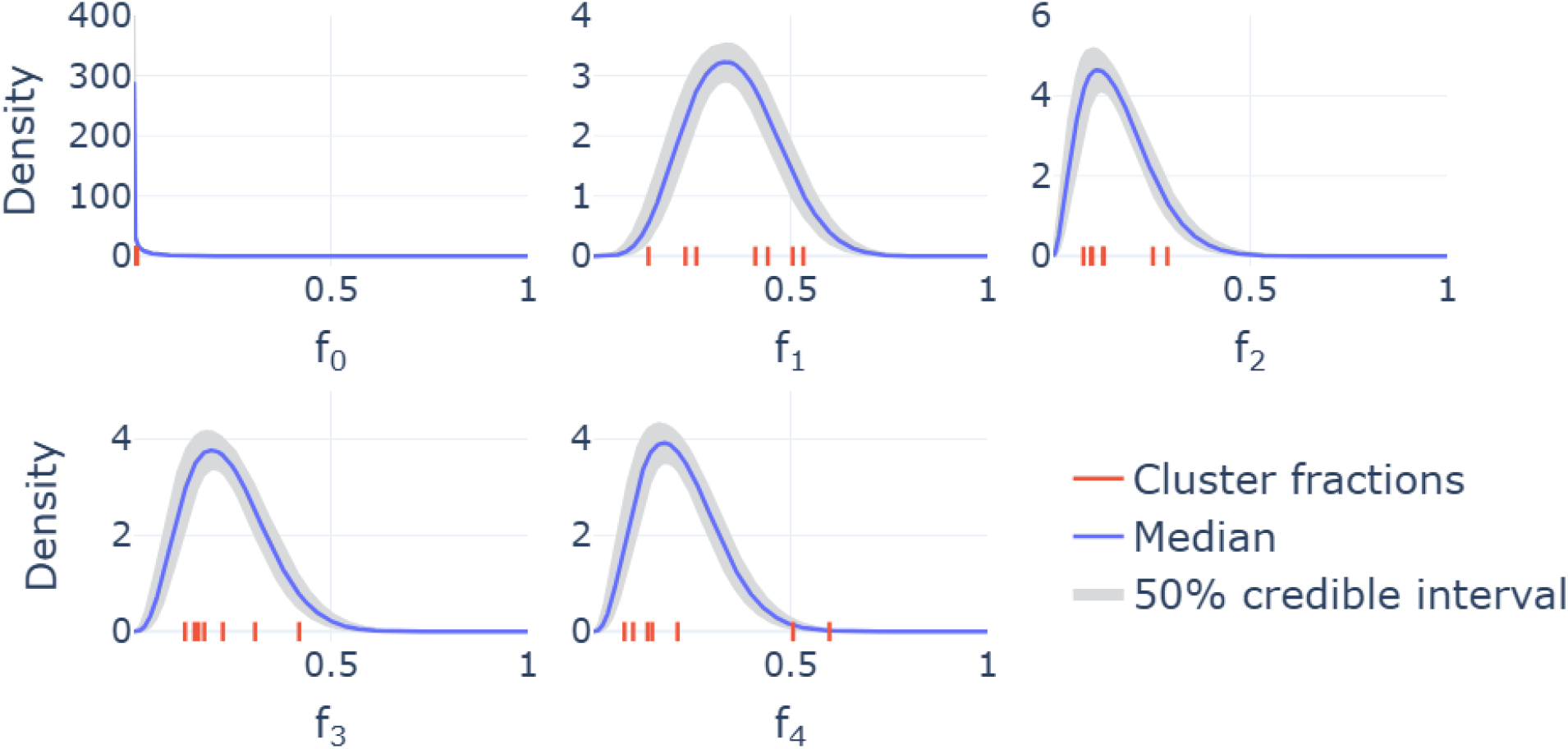
Marginal distributions describing the variability in cluster fractions across individuals. On each plot, the seven red ticks along the *x*-axis represent the cluster fractions for the seven individuals whose samples are included in the single-cell sequencing analysis. The median curves across these predicted densities are shown by solid lines, while the shaded 50% credible intervals indicate regions that half of these densities fall within.

**Figure 5.**
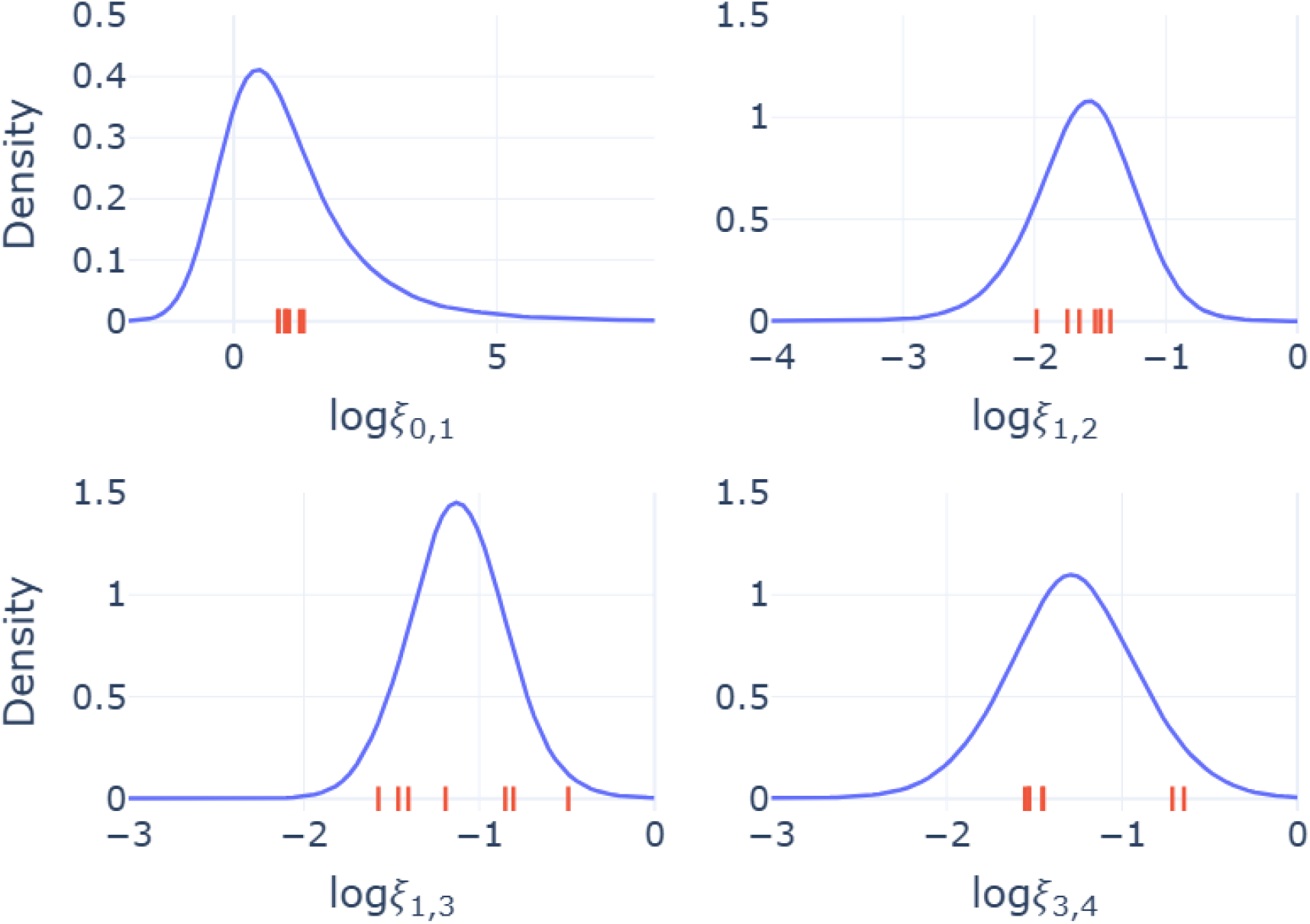
Probability distributions for the transition rates, *ξ*_*i,j*_ between state *i* and state *j*. The red ticks on the *x*-axes indicate the values of each transition rate for the seven individuals included in the single-cell sequencing analysis. The solid lines are model-predicted densities and provide an indication of the likely values transition rates take across a broader population. Logarithmic values are in base 10.

Median estimates from this combined framework suggest a rapid transition from precursor to immature neutrophils (*ξ*_0,1_=4.9 *h*^−1^), consistent with the low observed fraction of cells in cluster 0. Transition rates between subsequent states occur on slower but comparable time scales: (*ξ*_1,2_=0.025 *h*^−1^, *ξ*_1,3_=0.076 *h*^−1^, *ξ*_3,4_=0.055 *h*^−1^). From these estimates, the fraction of neutrophils that follow the IFN-responsive path (via cluster 2) can be calculated as: *ξ*_1,2_*/*(*ξ*_1,2_ + *ξ*_1,3_)=0.24. This suggests that approximately 24% of neutrophils transition into the IFN-responsive state, while the remaining 76% continue along the standard maturation pathway.

## Discussion

This study contributes to our understanding of neutrophil heterogeneity by introducing a novel mathematical framework to model transcriptional state dynamics in circulating neutrophils in adult healthy subjects. While mathematical models have been widely applied to study other immune cell populations, particularly T cells, relatively few efforts have addressed the transcriptional progression of neutrophils. Here, we leveraged single-cell RNA sequencing data from healthy individuals to identify five distinct transcriptional neutrophil states through clustering analysis. Building on previous work, and based on the expression of key marker genes, these clusters were interpreted to represent a progression from precursor cells to more mature states, with one cluster reflecting a type I IFN-responsive population [6, 15]

By integrating single-cell transcriptomic data with a system of ODEs, we provide a quantitative model for neutrophil state transitions. This model not only formalizes hypotheses about neutrophil maturation pathways but also enables the estimation of transition rates between transcriptional states. Despite a limited number of healthy subject samples, our statistical approach, based on Bayesian inference, allowed us to incorporate inter-individual variability and generate population-level estimates of state dynamics. This feature is especially valuable for extending insights beyond the specific individuals sampled and provides a framework that can scale with larger datasets of individuals affected by a number of disease states or physiological events [1, 3].

A key insight from the model is the fast transition rate from the precursor to immature state (*ξ*_0,1_), consistent with the low fraction of neutrophils observed in the precursor cluster. However, we note that the inferred values for *ξ*_0,1_ are highly sensitive to small variations in the precursor fraction (*f*_0_), which is estimated from a sparse population (≤1% of cells). Since *ξ*_0,1_ is approximately equal to *µ/f*_0_, small fluctuations in *f*_0_ can lead to large changes in *ξ*_0,1_. Thus, while the exact value of *ξ*_0,1_ may be uncertain, the main conclusion, that this transition occurs rapidly, remains robust. In future studies, it may be reasonable to exclude the precursor cluster and return to the four-cluster model identified in previous work, particularly if more robust estimates of *f*_0_ remain difficult to obtain [6].

We also assume a neutrophil clearance rate (*µ*) corresponding to a 12-hour half-life in circulation, consistent with established estimates. Since each transition rate in the model scales linearly with *µ*, uncertainty in *µ* propagates proportionally to the estimated rates. A +10% change in *µ*, for example, would result in a corresponding +10% change in all transition rate estimates. Moreover, the current statistical model assumes that all individuals are sampled from a single population. With a larger and more diverse dataset, it would be possible to incorporate covariates such as age, sex, or clinical status to refine these assumptions and better capture population-level variability.

Importantly, our model examines and assumes a steady-state neutrophil population (in the different states) for healthy individuals. Extending the framework to capture non-equilibrium dynamics, such as those occurring during acute infection, chronic inflammation, or drug treatments, would represent a valuable future direction. In addition, while our model is built on transcriptional data, integrating complementary data types (*e*.*g*., proteomics, epigenetics, functional assays, or tissue localization) would provide a more comprehensive view of neutrophil function and state transitions [11, 2]. We point out that our cluster interpretation is consistent with previous work [6], yet, the current study goes further by placing these clusters within a dynamic mathematical model. This framework provides not only a snapshot of neutrophil heterogeneity, but also insight into the timing and variability of transitions between states, offering a new dimension to understanding neutrophil maturation.

In summary, this study presents a foundational framework for modeling neutrophil transcriptional dynamics, addressing a significant gap in immunological modeling. Beyond improving our understanding of neutrophil behavior in healthy individuals, this approach lays the groundwork for investigating how neutrophil state dynamics are altered in disease contexts, including autoimmune and auto-inflammatory disorders, infections and metabolic diseases. Ultimately, the integration of mathematical modeling with high-dimensional single-cell data can inform both mechanistic understanding and experimental design, advancing the study of neutrophil biology and its role in immune system function.

## Acknowledgments

SP was supported by the NIH 2022 National Service DATA Scholarship. The study was also funded in part by the Intramural Research Program at NIAMS/NIH (ZIA AR-041199). The authors declare no conflict of interest.

This manuscript has been reviewed at Los Alamos National Laboratory and assigned report number LA-UR-24-27105.

